# Hyperspectral image analysis for classification of multiple infections in wheat

**DOI:** 10.1101/2025.07.16.664902

**Authors:** Manon Chossegros, Amelia Hubbard, Megan Burt, Richard J. Harrison, Charlotte F. Nellist, Nastasiya F. Grinberg

## Abstract

Plant diseases can cause heavy yield losses in arable crops resulting in major economic losses. Effective early disease recognition is paramount for modern large-scale farming. Since plants can be infected with multiple concurrent pathogens, it is important to be able to distinguish and identify each disease to ensure appropriate treatments can be applied. Hyperspectral imaging is a state-of-the art computer vision approach, which can improve plant disease classification, by capturing a wide range of wavelengths before symptoms become visible to the naked eye. Whilst a lot of work has been done applying the technique to identifying single infections, to our knowledge, it has not been used to analyse multiple concurrent infections which presents both practical and scientific challenges. In this study, we investigated three wheat pathogens (yellow rust, mildew and Septoria), cultivating co-occurring infections, resulting in a dataset of 1,447 hyperspectral images of single and double infections on wheat leaves. We used this dataset to train four disease classification algorithms (based on four neural network architectures: Inception and EfficientNet with either a 2D or 3D convolutional layer input). The highest accuracy was achieved by EfficientNet with a 2D convolution input with 81% overall classification accuracy, including a 72% accuracy for detecting a combined infection of yellow rust and mildew. Moreover, we found that hyperspectral signatures of a pathogen depended on whether another pathogen was present, raising interesting questions about co-existence of several pathogens on one plant host. Our work demonstrates that the application of hyperspectral imaging and deep learning is promising for classification of multiple infections in wheat, even with a relatively small training dataset, and opens opportunities for further research in this area. However, the limited number of Septoria and yellow rust + Septoria samples highlights the need for larger, more balanced datasets in future studies to further validate and extend our findings under field conditions.

## Introduction

Wheat (*Triticum aestivum*) is one of the world’s main food sources; globally, 790 million tonnes of wheat were produced in 2023 (1). Pathogens represent a significant threat to food security, with yellow rust (YR; *Puccinia striiformis* f. sp. *tritici*), mildew (*Blumeria graminis* f. sp. *tritici*) and Septoria (*Zymoseptoria tritici*) amongst the most common wheat foliar diseases in the United Kingdom. Collectively, they cause 15-30% of the annual wheat losses (2,3). These foliar pathogens are identified by their characteristic visual manifestations, which at a later stage of disease progression can be seen with the naked eye. All three pathogens cause a reduction in green leaf area and therefore limit the plant’s photosynthetic capability, leading to a reduction in yield. Yellow rust, also known as stripe rust, forms long yellow fine stripes on the leaves, composed of small pustules called uredia (4), mildew develops into white fluffy pustules on the surface of the plant, which are called conidiophores (5), whilst Septoria presents as irregular pale brown zones that can contain tiny dark fruiting bodies called pycnidia (2,6).

Today, disease recognition is mainly done ‘by eye’, which requires expertise from an assessor and can be error-prone (7,8). Computer vision for disease recognition has thus become increasingly appealing, mostly because it can detect changes invisible to the naked eye and automate the detection process (9). In the context of simultaneous infection, however, studies have shown that the computer vision can fail to label RGB images correctly (3). Hyperspectral images (HSI), on the other hand, are able to utilise a wider range of wavelengths, that go beyond the visible spectra, revealing changes in pigmentation or cell structure due to the presence of pathogens (10–12). The output is a 3D image called hypercube (13) (Figure 1a), with each pixel possessing a 2D spectral representation: reflectance of the pixel under different wavelengths (Figure 1b).

**Figure 1.**
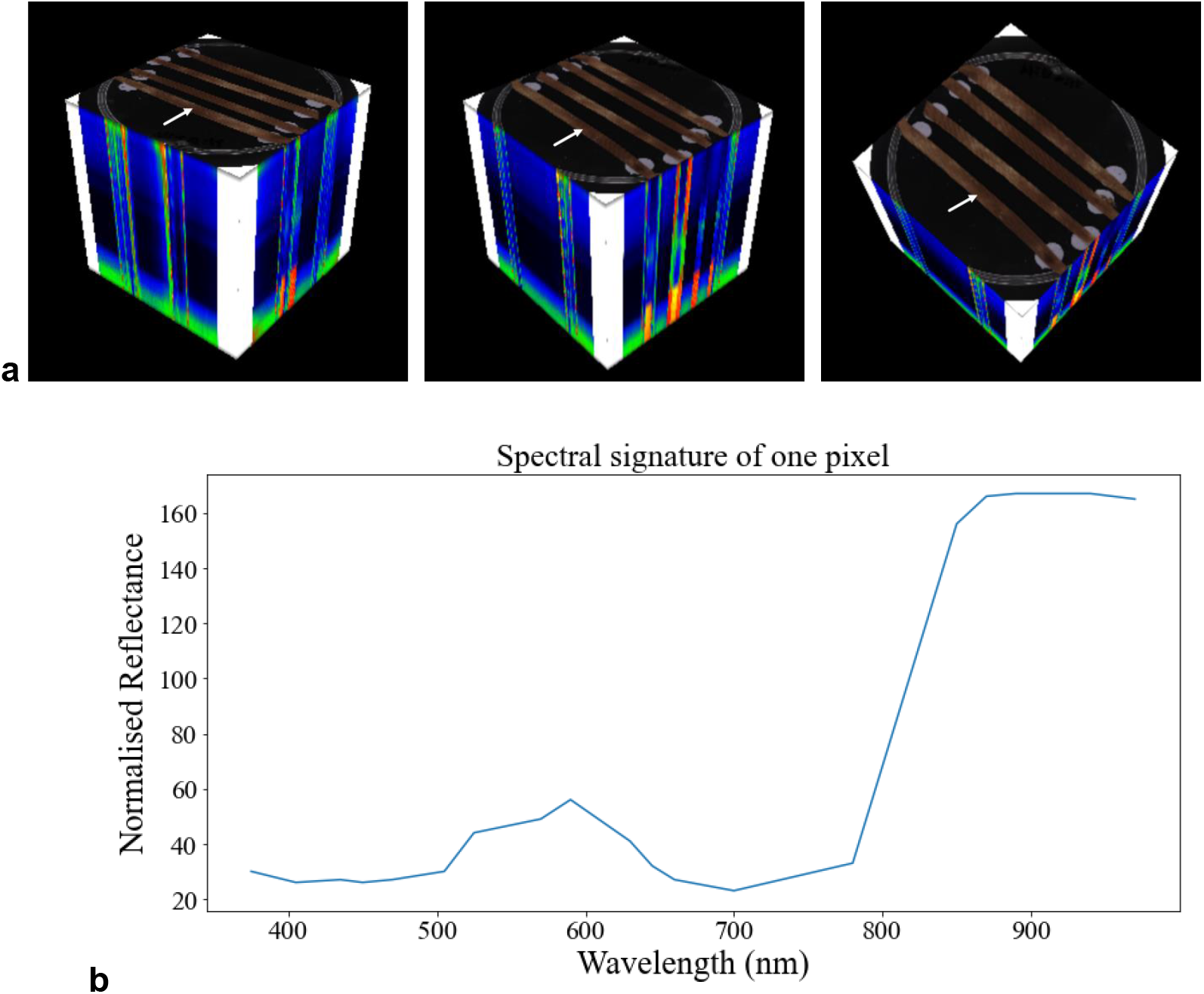
Example hyperspectral image of four wheat leaves a) a hypercube from three rotational angles, and b) representation of the spectrum of a single pixel (pointed to by the white arrow in a)).

Plant pathogens can cause disease in plants through various strategies of infection (8). Yellow rust and Septoria enter the plant via the stomata, a pore at the surface of the leaf responsible for gas exchange (14). After infection by Septoria, the disease stays in a latent asymptomatic phase for 14-28 days, before entering the necrotic phase, creating irregular pale brown zones (14). Yellow rust’s uredia typically appear after ten days. Mildew will show visible spores (conidium) after 5-10 days post infection (6).

In the field, however, diseases do not generally occur in isolation, and the presence of multiple diseases is common in plant disease epidemics (8). Two pathogens coexisting on the same plant are likely to interact, which can increase or decrease the virulence of the infection and potentially change expression of either or both diseases. These potential interactions between multiple pathogens can confound classifiers trained on single diseases leading to misdiagnoses, wrong treatment strategies, and ultimately to yield losses.

Coexistence of Septoria, yellow rust and mildew is complex due to differences in lifestyles. Both yellow rust and mildew are obligate biotrophs and thus require living tissue to survive, while Septoria is a hemibiotroph, initially starting off as a biotroph and switching to necrotrophy, feeding on dead host tissue (2). According to studies (6,15,16), when the combined infection is successful, the proportion of infected tissue is smaller than the sum of the effects by individual pathogens. Interestingly, (17) report that yellow rust’s pustules close to necrotic areas of Septoria appeared browner than their usual bright yellow orange, which could be a manifestation of the interaction between pathogens. It has previously been observed (17) that the combined infection of yellow rust and mildew reduced both the surface of infected areas and the spreading of the spores. This mutual virulence reduction is thought to be due to the high activity of plant defence mechanisms in the presence of both pathogens, and the scarcity of available nutrients the plants have to share.

Computer vision has been widely used for plant disease recognition using digital RGB images (18) and there has been a particular drive to use neural network (NN) architectures for this purpose (see reviews (19,20)). Moreover, deep learning approaches showed better results than traditional machine learning approaches like random forest and support vector machines (21). In (3), for example, a neural network was designed to label 2,414 images of wheat leaves into seven categories, including healthy and seedlings. Of the leaves, just 6% were infected with multiple pathogens. Although the algorithm achieved high accuracy classifying single diseases, it showed mixed results for classification of multiple co-infections. This could be due to insufficient numbers of samples with multiple co-infections as well as due to some co-occurring infections presenting very similarly. Hyperspectral imaging can potentially deal with the latter problem. Despite extensive application of HSI to classification of single infections (11,22,23), to our knowledge, no studies have addressed concurrent infections, which are common in the field and thus present unique biological and computational challenges.

Infection by various pathogens affects not only visible RGB wavelengths but also bands between 600 and 735 nm (nanometre), which correspond to chlorophyll absorption and thus reveal the photosynthesis capacity of the imaged parts of the plant (24), and wavelength regions of 750 nm and higher, characteristic of the water content of the leaf (25). Both the plant’s chlorophyll absorption and its capacity for photosynthesis can be affected by pathogen infection. Hence, hyperspectral images, through covering a whole range of non-RGB wavelengths, can provide complementary information related to presence of infection and disease progression. This can be particularly useful when multiple infections are present. In this study, we created a dataset of hyperspectral images of wheat leaves: uninoculated as well as infected by several individual pathogens and their pairwise combinations. We studied and compared spectral profiles of single and concurrent infections and trained a selection of DL models to accurately classify the infections and their combinations. In particular, this study makes three contributions: (i) creation of the first hyperspectral dataset of concurrent wheat infections, (ii) application of Inception (26) and EfficientNet (27) with both 2D and 3D convolutions, assessing relative importance of spatial texture and spectral continuity, (iii) demonstration that concurrent infections (e.g. YR and Septoria) exhibit emergent spectral features resulting from interactions that are not simply the sum of the individual disease profiles.

## Materials and Methods

### Preparation of plant samples

The yellow rust, mildew and Septoria susceptible wheat variety ‘Vuka’ (28) was sown in 7.5 cm diameter pots and grown in a disease-free growth room at 17/11’ °C for 16/8 hours in the light/dark, for 10 days. Day of inoculation (Table 1) is counted from sowing of the wheat seeds. Plants were then moved to another growth room for inoculations, day and night temperatures were tailored to the pathogen, to maximise disease expression and are detailed in Table 1. Conditions were adjusted for multiple infections to synchronise symptom development.

**Table 1.** Summary of the inoculation conditions of wheat plants depending on pathogen.

| Disease | Day of inoculation | Day/night temperature (°C) |
| --- | --- | --- |
| Yellow rust | 10 | 18/11 |
| Mildew | 10 | 18/11 |
| Septoria | 10 | 22/12 |
| Yellow rust + mildew | 10 (yellow rust)<br>13 (mildew) | 18/11 |
| Yellow rust + Septoria | 10 (septoria)<br>17 (yellow rust) | 22/12 |

The yellow rust isolate used in this study was isolate WYR 19/215 provided by the United Kingdom Cereal Pathogen Virulence Survey (UKCPVS). Desiccated urediniospores were put into a suspension with hydrofluoroether solution and sprayed over the leaves using a compressor pump. To perform mildew inoculations, spores from infected leaves (NIAB 21-001 mildew isolate) were blown into settling towers placed over the healthy pots. Septoria inoculum R13 and R16 were harvested after one week from YPD (yeast extract peptone dextrose agar) plates. Spore concentration was then determined with a haemocytometer and diluted to 5 x 10^5^ spores per mL. A total of 10 mL of this solution was sprayed onto each pot.

For multiple infections, inoculations were planned so that symptoms of each disease were at their peak at approximately the same time. For the YR + mildew combination, yellow rust and mildew inoculations were performed at 10 and 13 days, respectively. For the YR + Septoria combination, yellow rust and Septoria inoculations were performed at 10 and 17 days, respectively.

Additional leaves expressing septoria symptoms were collected from the field and included in the analysis.

### Imaging of leaves

Four wheat leaves were placed adaxial side up, in 9 cm Petri dishes, affixed by Blu Tack. Care was taken to avoid damaging the leaves and thus affecting the symptom expression. Leaves exhibiting symptoms visible by eye were chosen to be part of the training dataset. These leaves were imaged with a lab-based multispectral imaging system, the VideometerLab 4 camera (see full specifications at (29)). Before the acquisition of images, the camera was calibrated using the measurement of a white and a black hyperspectral image. The final image was a hypercube of size 2,192 pixels x 2,192 pixels x 19 with the 19 wavelengths covering the 375-970 nm range (Figure 2). A total of 1,447 images were taken (Table 2).

**Table 2.** Final number of images generated for each treatment.

| Treatment | No. of imaged leaves |
| --- | --- |
| Uninoculated | 359 |
| Yellow rust | 275 |
| Mildew | 362 |
| Septoria | 36 |
| YR + mildew | 376 |
| YR + Septoria | 39 |
| **Total** | **1,447** |

**Figure 2.**
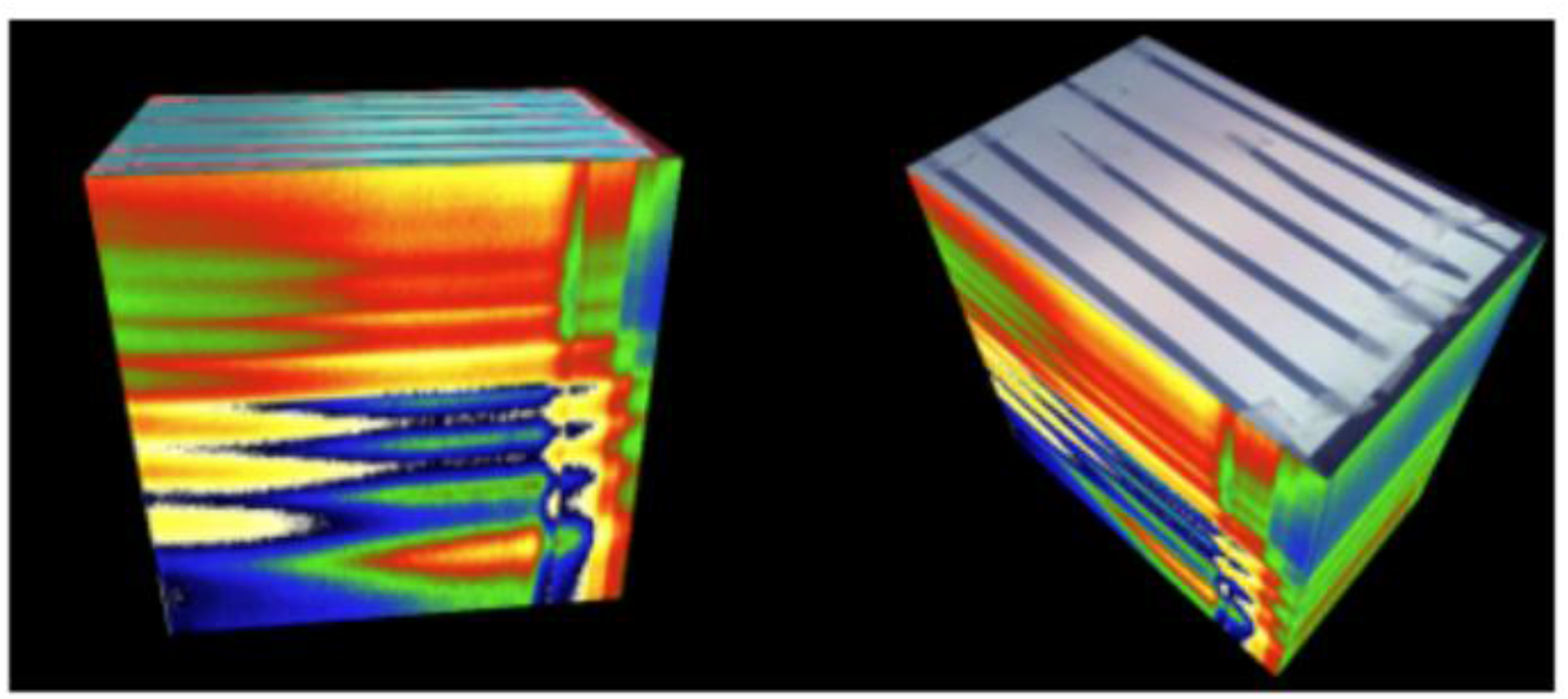
A sample hypercube (from two rotational angles).

### Classification Models and defining class labels

The HSI dataset was analysed using two ImageNet-pretrained convolutional neural network (CNN) models: Inception-V3 (26) and EfficientNet-B0 (27), each with a choice of either a 2D or 3D convolutional layer as the first step, resulting in four possible architectures.

Instead of a multilabel classification setting, where samples infected with multiple diseases could belong to more than one class, the images were labelled with six classes: Uninoculated, Septoria, mildew, YR, YR + mildew, YR + Septoria (there were no Septoria + mildew samples). This way each pairwise combination of infections corresponded to its own label with potential interactions between different diseases taken into account.

### Image pre-processing and preparation of the training dataset

The specific parts of the image that were used as input samples were carefully selected, finding a balance between reducing the large size of the hyperspectral cube and keeping as much information as possible. Images were cropped to keep only the leaf area (Supplementary Fig. 1), and background was removed using thresholding followed by erosion and dilation in order to reduce noise (see e.g. (12)). Finally, images were cropped to keep only one leaf per image (Supplementary Fig. 2). In order to reduce the noise in the spectral dimension due to artefacts or abnormally bright bands, Savitzky-Golay smoothing was applied as described in (30).

In order to increase the size of the training data, the original images were supplemented with their distorted versions, vertical flips and random rotations (see e.g. for a similar treatment (31); Supplementary Fig. 3). This not only increased the size of the training data but also increased the variety of available images and made the algorithms more robust to image distortions.

### Band selection

To better understand the biological process of the disease as well as to select the best bands for dimensionality reduction, the importance of each wavelength was determined. For each disease and combination of diseases, 350 infected regions of interest (ROI) of size 30 x 30 pixels^2^ were selected. Additionally, 350 ROI of the same size were selected on the uninoculated leaves. ROIs were extracted by visually determining symptomatic zones of the leaves. These zones were then delineated on the hyperspectral images and subdivided into squares based on their pixel coordinates using Python (CV2). For each of the 19 bands, average reflectance of each ROI in symptomatic leaves was measured and compared to the uninoculated controls via an independent samples *t*-test under the assumption of equal variances (as in (32)). The significance level was set to 5% and the Bonferroni correction (within each class) was applied to account for multiple testing.

To reduce the dimensionality of the problem and to alleviate the computational burden, the top four most significant (as per test statistic values) wavelengths for each of the five disease categories (Supplementary Table 1) were selected and their union used for training our models. Additionally, 570 nm and 590 nm wavelengths were also used because they were representative of the green and yellow colour changes and could visually discriminate Septoria from yellow rust. Hence, the wavelengths (in nm) used for model training were: 570, 590, 645, 660, 700, 780, 870, 890, 940, 970, with the final size of the images being, after pre-processing, 1,350 x 500 x 10 pixels^3^.

### Training and validation of the models

On our dataset, we trained and validated predictive accuracy of the following four architectures: 2D convolutional layer + Inception-V3 (2D Inception), 3D convolutional layer + Inception-V3 (3D Inception), 2D convolutional layer + EfficientNet-B0 (2D EfficientNet), and 3D convolutional layer + EfficientNet-B0 (3D EfficientNet).

Each model consisted of the pretrained architecture of EfficientNet-B0 or Inception-V3 to which a first convolutional layer was added. This layer was either a 2D or 3D filter and contained 10 channels corresponding to the 10 wavelengths of the input hypercube. The convolutional layer then fed the resulting 3-channel image to EfficientNet or Inception model. Finally, a last linear layer mapped the output of the pretrained backbone to 6 disease classes.

For each architecture, we performed 3-fold cross validation whereby during each iteration, the model was trained on the two thirds of the dataset and tested on the remaining third. Thus, predictions were obtained for every sample in the dataset. Models were trained on GPU with stochastic gradient descent using the following hyperparameters: *learning rate*=0.1 and *momentum*=0.9. We optimized the models through *CrossEntropyLoss* with a batch size of 6.

The performances of each network were evaluated according to the following metrics:

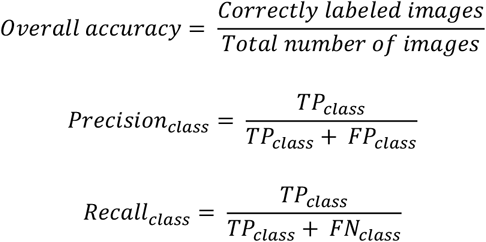

where *TP*_*class*_, *FP*_*class*_ and *FN*_*class*_ denote true positives, false positives and false negatives for a given class, respectively. Precision and recall were calculated per class whilst accuracy is an overall assessment of the algorithm’s performance. Note that recall, proportion of correctly classified instances for a given class, is essentially class accuracy, relating to the algorithm’s ability to identify *all* samples of a given class. Precision, meanwhile, is proportion of instances classified as the current class, that is correct. It is high when the algorithm’s labels for a given class are accurate, even if not all samples of this class are identified.

## Results

This study created a dataset of 1,447 hyperspectral images of uninoculated (Figure 3a), single and double infections of wheat leaves (Table 2), enabling the classification of multiple co-infections using spectral imaging. Symptoms of yellow rust (Figure 3b) and mildew (Figure 3d) were consistent and stable, whilst Septoria symptoms were highly variable and presented challenges within the timeframe of the experiment. The double infection of mildew + Septoria could not be imaged and only a few leaves presented symptoms of Septoria, and yellow rust + Septoria, which was at a very advanced stage. To counter the lack of Septoria images, 16 additional leaves were also collected from the field (Figure 3c). They had the main advantage of showing good symptoms of Septoria, but they were not the same age or of the same variety of wheat as our laboratory-grown wheat.

**Figure 3.**
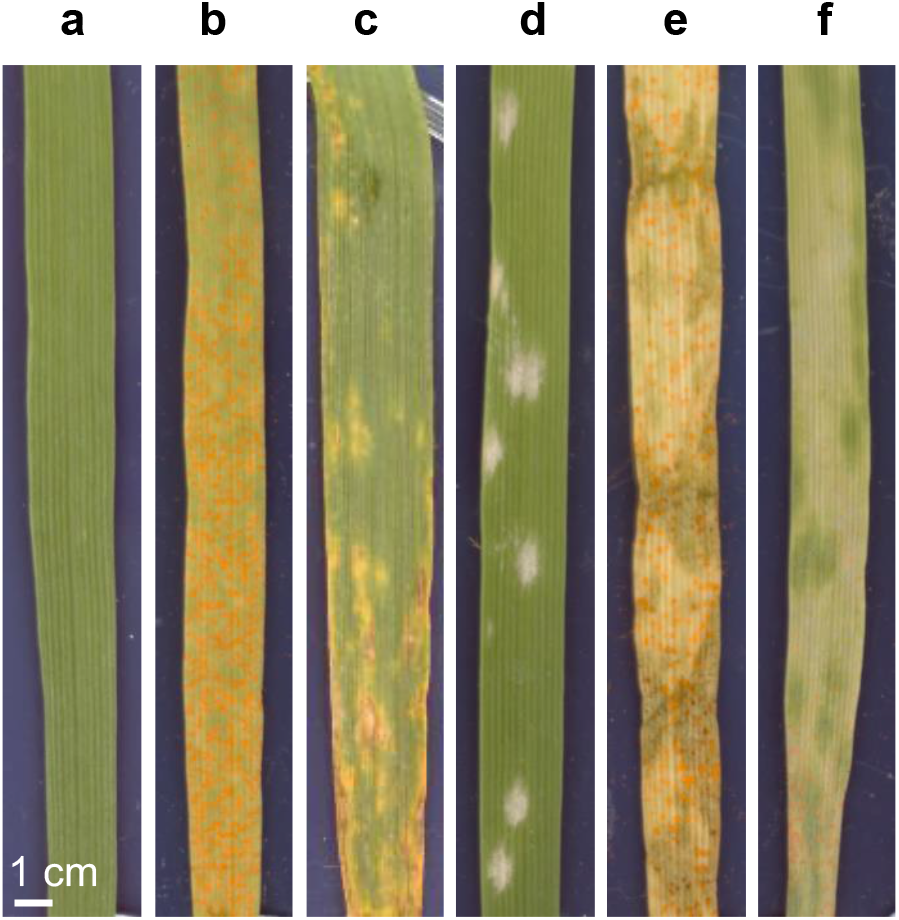
Example RGB images of symptoms of the wheat foliar diseases and their combinations investigated in this study, a) uninoculated, b) yellow rust (YR), c) Septoria, d) mildew, e) yellow rust + Septoria, f) yellow rust + mildew.

The expected visible manifestations of each disease were observed, urediniospores for yellow rust, conidiophores for mildew, and necrotic regions with pycnidia for Septoria (examples in Figure 3). The combination of multiple diseases exhibited both pathogen symptoms (Figure 3 e) and f)). It was also observed in a combined YR + Septoria infection, yellow rust pustules appear darker in colour than the bright orange of a single infection and the combined infection of yellow rust and mildew exhibited reduced symptoms from both diseases compared to single infections.

The average reflectance for each disease and pairs of diseases, as well as the uninoculated leaves, at each of the 19 wavelengths were plotted (Figure 4).

**Figure 4.**
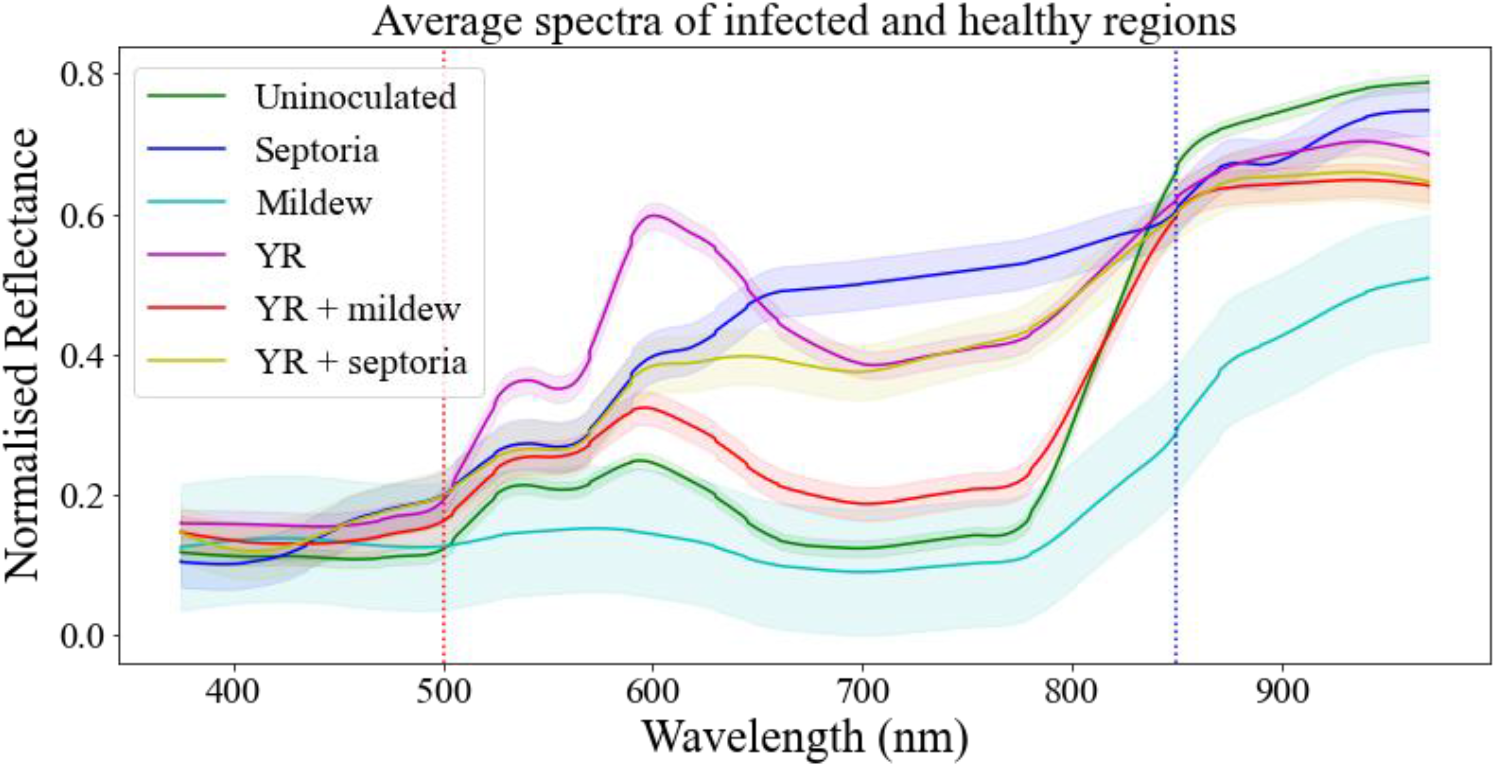
Difference in average reflectance between uninoculated and infected leaf surfaces. Bold lines represent average reflectance over 350 uninoculated (green) and 350 infected ROI for every single pathogen and the combined infections. Shaded envelopes designate +/-standard deviation for each wavelength. Vertical dotted red and blue lines designate values of interest 500 nm and 580 nm, respectively. Normalised reflectance was obtained by dividing the reflectance values by 255 to achieve the [0,1] scale.

There were some notable differences and similarities between the graphs for the five disease classes. A series of *t-*tests comparing reflectance for each band between each disease and Uninoculated leaves showed that the differences we can observe in the graph are statistically significant (with the exception of mildew at 505 nm; Supplementary Table 2). The largest differences between the signatures were observed for wavelengths above 500 nm and below 850 nm (apart from mildew; Figure 4).

Notably, we see that the spectral signature of the yellow rust and Septoria combination does not look like spectral signatures of either of the individual infections suggesting that there is an interaction between co-occurring diseases. Reflectance for the double infection of Septoria + YR (yellow line) seems to align closely with reflectance for Septoria (blue line) until around 600 nm and with reflectance for yellow rust (purple line) after around 700 nm with the 600-700 nm range looking distinctly different from both individual diseases. Moreover, the 570 nm and 590 nm wavelengths were particularly helpful in discriminating between the two co-occurring infections (Figure 5). Similarly, the yellow rust and mildew combination presents a different spectral profile (red line) to the two individual diseases: yellow rust (purple line) differs significantly from the combination in the 500-800 nm range while mildew (turquoise line) diverges from the combination after 500 nm. As noted, above, combined YR + Septoria infection reduces symptoms of both infections and this is reflected in the reflectance graph with the double infection line (yellow line) resembling more in profile that of the uninoculated/healthy leaves (green line) than of yellow rust (purple line) or mildew (turquoise line). However, in both cases of double infections, YR + Septoria and YR + mildew, the spectral signature of the combination resembles more Septoria and mildew, respectively, rather than yellow rust (Supplementary Table 2). This might indicate that yellow rust is dominated by its co-infections or that the conditions were more optimal for the other pathogens.

**Figure 5.**
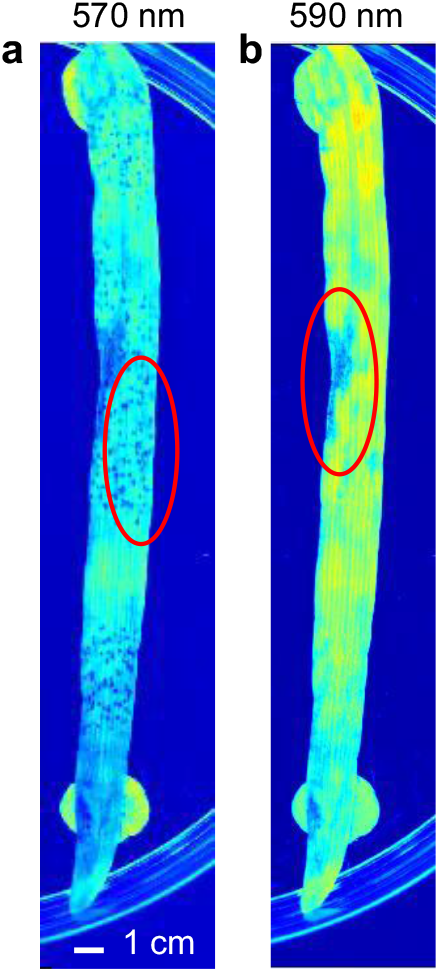
Discriminatory wavelengths: visualisation of Septoria and yellow rust at different wavelengths, with respective symptoms circled, a) emphasis on YR at 570 nm b) emphasis on Septoria at 590 nm.

Out of the four architectures investigated, the highest overall accuracy was achieved by EfficientNet-B0 with a 2D convolutional layer. It showed the highest precision and recall for every class apart from Septoria (Table 3). The better performances of EfficientNet were probably due to its ability to capture information both in resolution and in width, which was particularly relevant for spectral images. Furthermore, EfficientNet uses feature fusion which may have helped using both low-level and high-level features effectively.

**Table 3.**
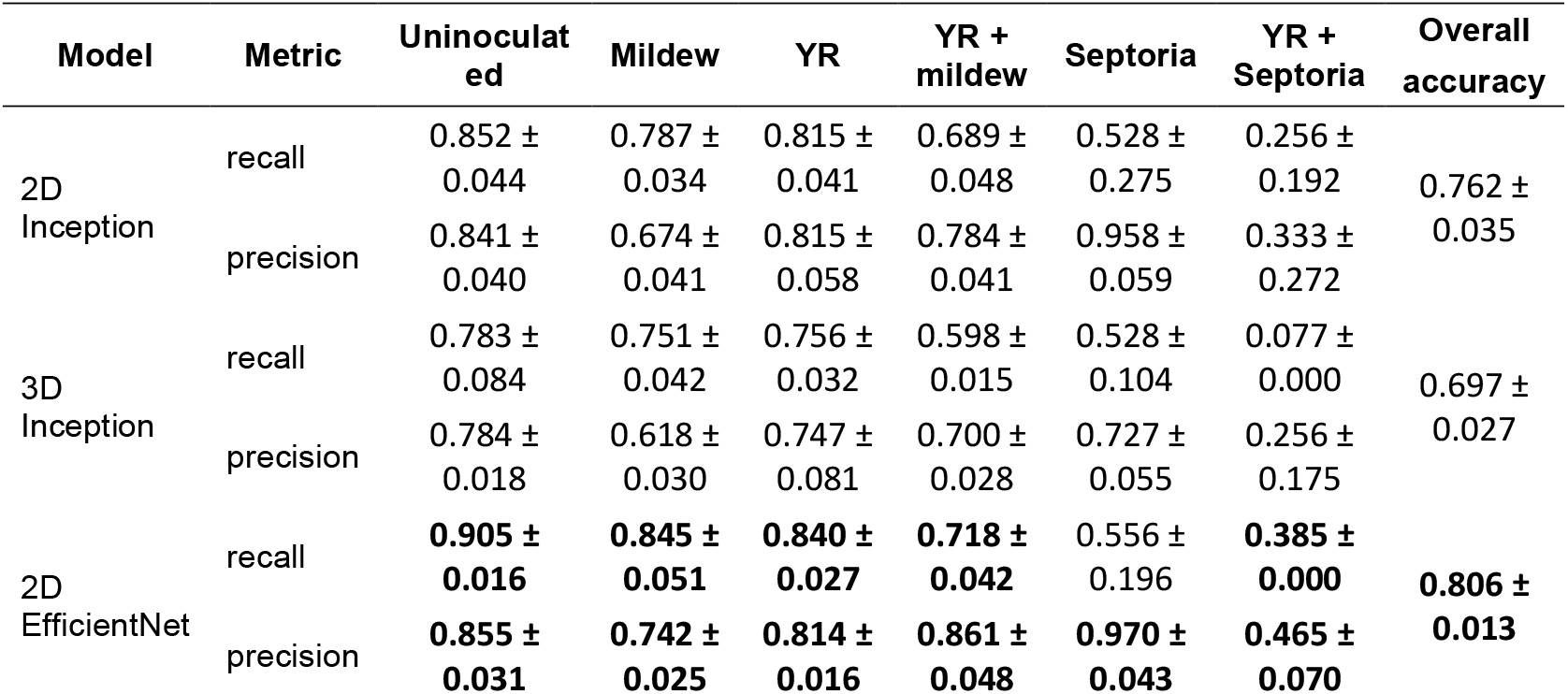
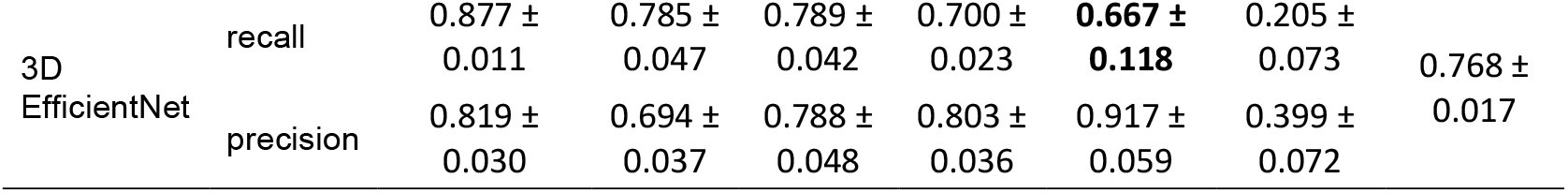
Overall accuracy and per class precision and recall for the four architectures. For each class column highest precision and recall are in bold; highest overall accuracy for the four methods is in bold.

Looking closer at the prediction accuracy per type of infection for the 2D EfficientNet model (Figure 6), note that it successfully predicted most instances of single infections for mildew (84.5%) and yellow rust (84%) as well as for the YR + mildew combination (71.8%), the latter result never having been described before. Accuracy for Septoria and YR + Septoria classes was substantially lower, likely due to the reduced number of imaged leaves (36 Septoria and 39 Septoria + YR). Moreover, the YR + Septoria class suffered not only from low sample numbers but also from biological variability stemming from Septoria’s variable latent period which led to asynchronous infection peaks. This produced heterogeneous symptom expression reducing the models’ ability to learn discriminating features for the combined infection class. Additionally, the Septoria class contained both lab grown and field samples creating intraclass variability due to differing growth conditions (e.g. light, temperature, etc.) and making generalisation harder. Thus, the low performance on Septoria and YR + Septoria classes reflects sample scarcity and quality, and biological variability rather than fundamental limitations of the CNN architectures.

**Figure 6.**
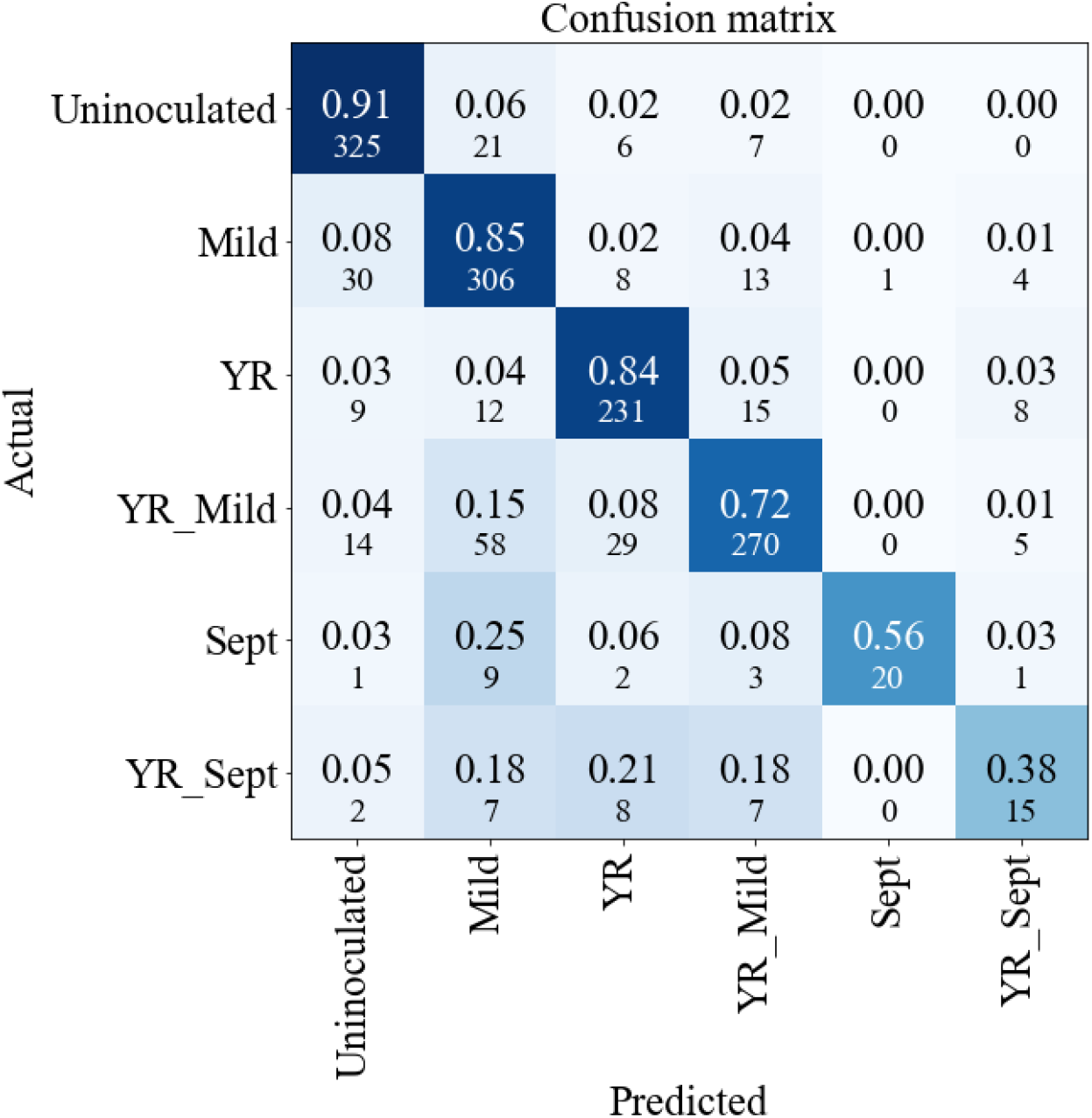
Confusion Matrix for 2D EfficientNet. Observed classes are in rows and predicted classes are in columns. The larger number in *ij*^*th*^cell is the proportion of class i predicted as class j with the smaller number being the corresponding number of samples.

Interestingly, in the case of multiple diseases, when an incorrect prediction was made, 86.7% of the cases for YR + mildew and 62.5% cases for YR + Septoria, the predicted label contained one of the pathogens present on the leaf (this can easily be calculated from the confusion matrix in Fig. 6).

## Discussion

Although ubiquitous, multiple co-occurring infections have not been given nearly as much attention in the computer vision community as they deserve. There is, to our knowledge, no studies applying hyperspectral imaging to double infections in wheat (or any other crop). In this study, we have attempted to bridge this gap by demonstrating the utility of a combination of deep learning approaches and hyperspectral imaging for identifying single and double infections in wheat. We have also highlighted potential difficulties that arise in this type of analysis.

Firstly, for lab-grown samples, an important technical area to explore was optimising protocols for successfully cultivating multiple infections. Despite carefully timing each inoculation, in the case of double infections, problems were still encountered producing viable YR + Septoria samples. The yellow colour of leaves infected by YR + Septoria came from non-simultaneous peaks of infection, resulting in yellow rust being at a very advanced stage when Septoria appeared. Septoria’s latent period can vary by up to two weeks, depending on the environmental conditions and variety. As a consequence, the two Septoria classes together accounted for 75 leaves, representing only 5.2% of the 1,447 samples. Expanding the work to include additional cultivars with varying degrees of susceptibility to the three different diseases would increase the robustness of the distinguishing wavelengths observed. In future work, *in silico* approaches could also be used to mitigate data scarcity and class imbalance. Options include oversampling and augmenting the training dataset with synthetically generated images. Generative approaches such as GANs (33) and, recently, diffusion models (34) have been shown to be useful for generating realistic looking plant disease images and might offer a way to expand underrepresented classes where data generation is experimentally laborious.

However, to increase the practical benefits from this study, conducted under controlled greenhouse conditions, and move closer to field deployment, there is a need to expand the collection of training samples to include field grown plants and plants at differing stages of infection. This would help to study co-evolution of concurrent infections by inspecting changes in their spectral signatures over time. Moreover, this would address a challenge presented by different diseases and their combinations exhibiting similar or overlapping spectral profiles to each other and to healthy leaves, especially at the early and intermediate stages of infection (35,36), making discrimination difficult. Incorporating a wider range of samples would shed more light on the dynamics of the interactions of concurrent infections on a host and help to develop techniques for early and severity detection of diseases. Additionally, this approach would help models generalise to environmental variability (illumination, background reflectance, canopy architecture) present under real life conditions. A model trained on a dataset complemented with real-life examples of single and concurrent infections would transform from a prototype to a practical tool, initially in a handheld device validated under field conditions, and later as hyperspectral sensors mounted on farm equipment (e.g. a tractor), or, potentially, an unmanned aerial vehicle (e.g. a drone), something that has already been done with single-infection HSI-based classifiers (37–39).

Plants are regularly exposed to co-infections by multiple pathogens under field conditions. Local environmental conditions, such as temperature, humidity and crop management practices strongly influence the prevalence and severity of these diseases. Controlling several diseases simultaneously presents a considerable challenge, since individual pathogens differ in their infection strategies and impact on yield and quality. Integrated approaches combine resistant cultivars, optimised agronomic practices and targeted chemical control, which needs to be based on accurate identification of diseases before they cause an epidemic. Ultimately, farmers must balance the suppression of multiple diseases with the need to maintain yield and profitability, highlighting the need for broad strategies that address multi-disease scenarios rather than treating pathogens in isolation.

Our analysis of the collected images revealed that the spectral signature of leaves infected with multiple diseases differed from the spectral signatures of the individual infections. This is something that can be noticed with the naked eye, which has been previously described in the literature (15, 40) and is now additionally confirmed by HS imaging. Presence in the training dataset, of samples infected with multiple diseases made our task of identifying them a multilabel classification consideration, where each sample can belong to more than one class. However, using the same class label for a disease whether it is the only infection affecting a leaf or one of a tandem of two infections, would ignore the fact that disease presentation might depend on any possible co-infections (e.g. pustule colour change). Treating each combination of infections as a separate class circumvented this problem as then any features characteristic of this combination were attributed to the combined class. The appearance of new spectral features in the case of combined diseases demonstrated that coexisting pathogens interact and thus behave differently when infecting a host together. This suggests that interaction between concurrent diseases potentially lead to changes in the molecular processes underlying the separate infections. Studying and understanding these biological processes, with the help of HSI, could lead to development of more efficient treatments and preventative measures for the agriculturally important crops. However, one-hot labelling also had drawbacks as it did not allow transfer of knowledge from single disease cases to co-infections, and it could be confused by differences in pathogen prevalence (e.g., a leaf showing mostly YR symptoms with a few Mildew spots). Future work could explore architectures that combine both approaches, e.g. simultaneously learning single disease patterns as well as modelling interactions between infections, possibly using an attention mechanism to capture interaction effects (41), or a dual-branch approach (42).

The choice of the model is of utmost importance for the classification results, and the literature reveals how critical this decision can be. For instance, one meta-study (21) compared a range of classifiers for crop disease detection and classification and found that CNN models consistently outperformed traditional ML approaches such as random forest and SVM. Additionally, CNN performance was sensitive to the choice of architecture and training strategy. Furthermore, image analysis, especially on smaller datasets, can be considerably improved by using models that have been pretrained. Hence, we chose to analyse our HSI dataset using two ImageNet-pretrained (43) CNN models: Inception-V3 (26) and EfficientNet-B0 (27), each with a choice of either a 2D or 3D convolutional layer as the first step, resulting in four possible architectures. In image analysis, CNNs involve convolution layers that pass an image through a 2D kernel (filter) (resulting in 2D-CNN), determining whether a local pattern is present in the image and thus deducing what sort of pixel arrangements are characteristic of the object class. However, to effectively use the wealth of information provided by HS images, a kernel would need to extract features spanning both spectral and spatial dimensions—a 3D kernel is needed, resulting in a 3D-CNN. It has been demonstrated in (32) that 3D-CNN performed better than 2D-CNN for the classification of grapevine disease, but higher complexity of the model comes at a computational cost and might suffer from overfitting. Inception and EfficientNet architectures both address the problem of achieving deeper networks whilst keeping dimensionality of the model manageable. More specifically, EfficientNet scales the width of the image, thus efficiently dealing with the spectral dimension and Inception uses small size convolutional filters to capture information at low cost. This made both models attractive choices for the described study. Note that for both algorithms, Inception and EfficientNet, a version with a 2D convolutional input layer performed better than a version with a 3D convolutional layer. Intuitively, one expects a 3D input layer to perform better than 2D layer due to its ability to detect characteristic patterns both in the spatial and spectral dimensions. The fact that the opposite turned out to be true in practice indicates the importance of patterns across neighbouring pixels, i.e. 2D texture, compared to the patterns across neighbouring wavelengths, i.e. the additional 3^rd^ dimension. Indeed, previously, several studies have highlighted the importance of texture features in plant disease recognition: for example, addition of texture features to hyperspectral data significantly improved accuracy of detection of yellow rust in wheat (22), and of early blight on eggplant leaves (44). It is likely that a 2D filter captures some of the texture information without it being included in the model explicitly. Moreover, relatively low performance of the 3D models compared to the 2D variants might be explained by the limited number of spectral bands (19) available from our imaging device compared to many other HSI studies which often use hundreds of bands (12,22,30). Explicit integration of texture and hyperspectral features for classification of co-occurring infections is a promising direction for future work.

Additionally, to better understand which spatial or spectral patterns drive the models’ decisions, and in particular identification of each disease or combination of diseases, future work could employ interpretability methods. Among these, saliency maps such as Gradient-Weighted Class Activation Mapping (Grad-CAM; (45)) can highlight pixels relevant to the prediction. In a multilabel setup, they might also help distinguish important regions in co-infected samples.

Whilst the four DL architectures we employed showed overall good accuracies, further avenues of research should include exploring the use of other DL approaches that may perhaps work better with smaller datasets (producing HS images of plant infections is laborious and DL methods are data hungry) such as Siamese (46) or relation networks (47). Additionally, our observation that 2D-CNN architectures outperformed the 3D-CNN ones highlights the importance of the leaf texture for detecting and identifying infections. Further exploration of this phenomenon and potentially explicit inclusion of texture features as inputs to classification algorithms (22) could prove to be fruitful.

Overall, we believe we have made an important first step into studying the application of HS imaging and DL to concurrent diseases in crops (with wheat as our study species). We devised a protocol for growing and inoculating wheat leaves with single and double infections and assembled a dataset of around 1,400 hyperspectral images. We demonstrated that using even our relatively modest dataset for training, high classification accuracies can be achieved by various CNN-based algorithms. Our work presents several directions for further research, including supplementing the training dataset with field samples with both single and multiple infections, the use of different classification algorithms, the use of alternative labelling, further study of the role of texture in classification of multiple infections and investigating interactions between concurrent infections using the HS imaging technology.

## Availability of Code

All analyses were conducted in Python with DL models deployed using Keras (48). Code for training, deploying and testing the models can be found at https://github.com/mc2295/hyperspectralplants.

## Acknowledgements

We thank the United Kingdom Cereal Pathogen Virulence Survey (UKCPVS; funded by Agriculture and Horticulture Development Board, AHDB and the Animal and Plant Health Agency, APHA) for providing the yellow rust isolate, WYR 19/215, used in the study. The authors gratefully acknowledge the support and advice on inoculation procedures from Ms Eda Naska and Ms Sarah Wilderspin, as well as Prof. Ji Zhou for providing assistance and support with the collection of images and for useful discussions.

## Author Contributions

R.J.R., C.F.N. and N.F.G. conceived the project and supervised all ongoing work. M.C., A.H. and M.B. carried out laboratory growth experiments. M.C. performed imaging and all numerical analysis and wrote the manuscript with contributions from N.F.G. and C.F.N. All authors reviewed the manuscript.

## Ethics Declarations

### Ethics approval and consent to participate

Not applicable.

### Consent for publication

Not applicable.

## Supplementary Figures and Tables

**Supplementary Figure 1.**
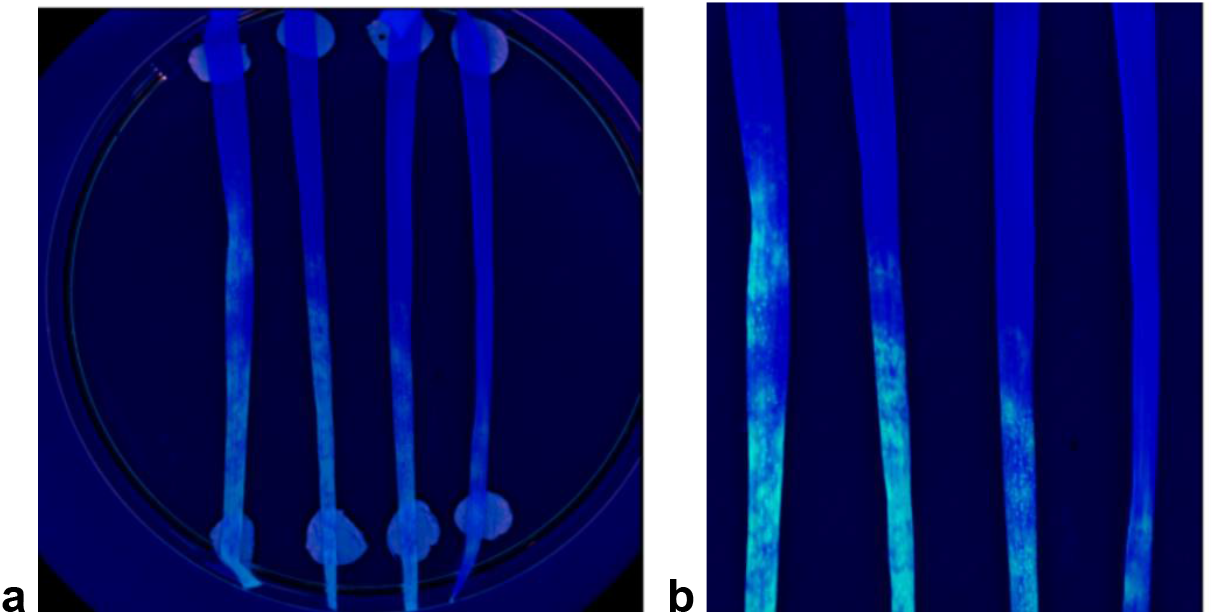
Cropping and preprocessing of the image: a) the original image, b) image after noise reduction and cropping.

**Supplementary Figure 2.**
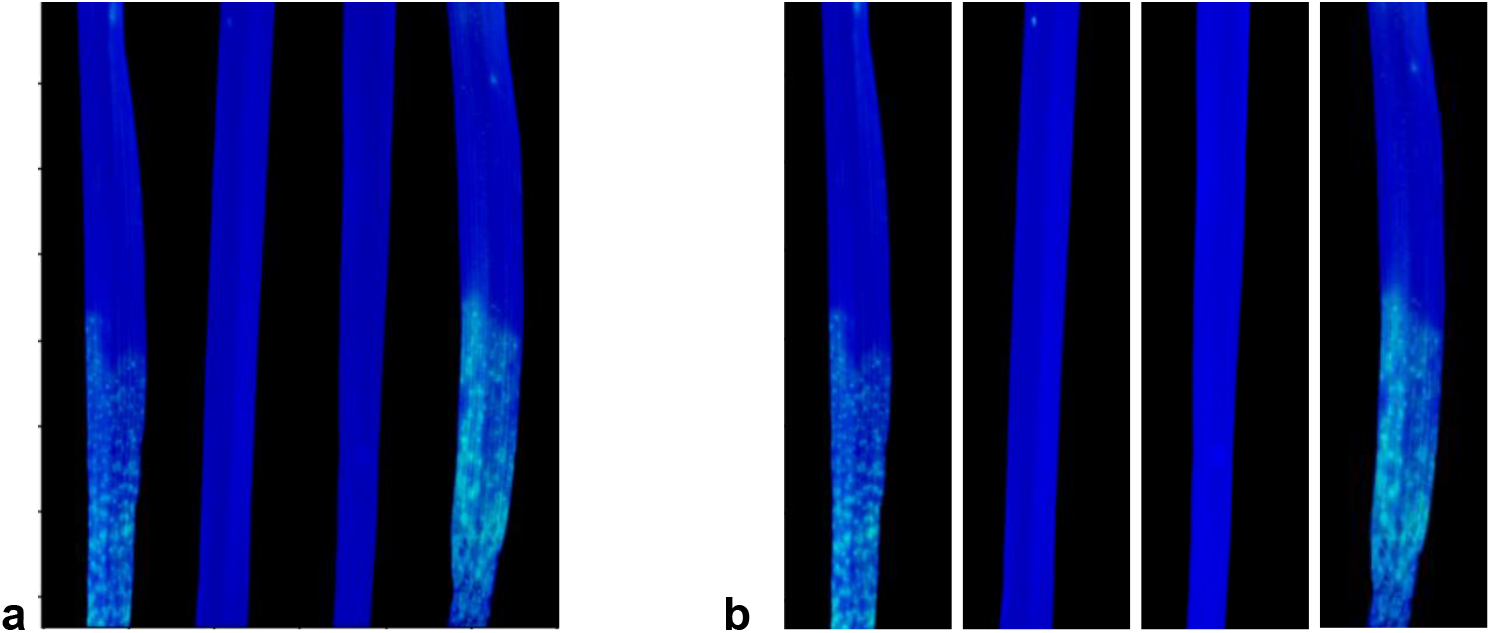
Final leaf separation: a) initial image b) separation into individual leaves -- input samples.

**Supplementary Figure 3.**
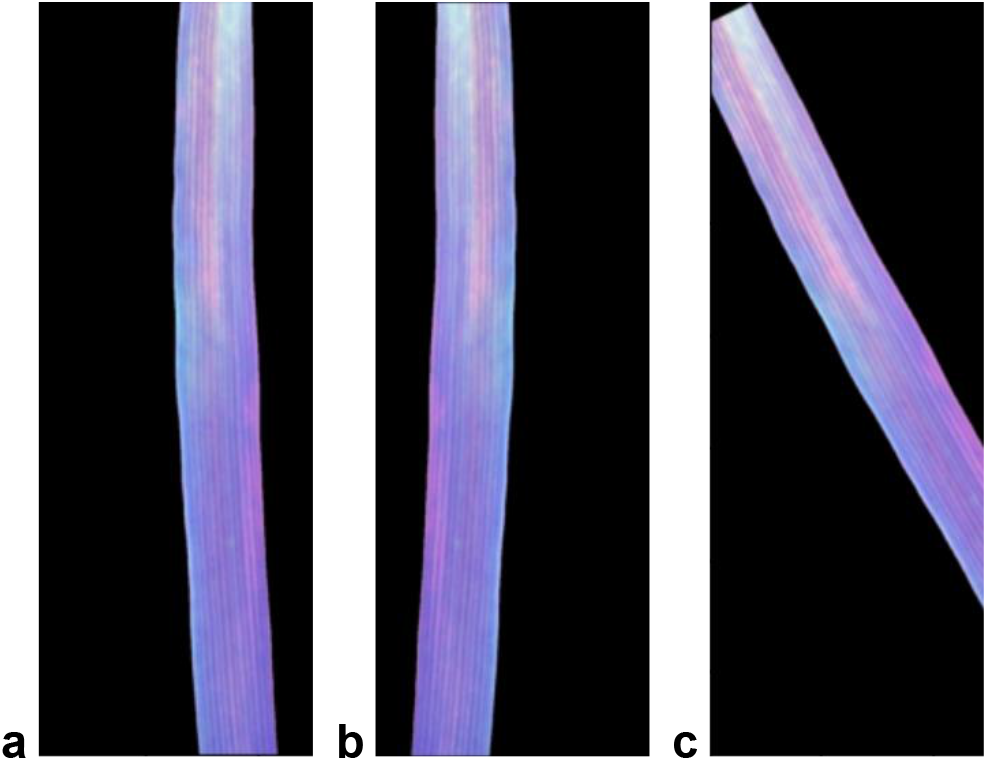
Data augmentation: a) original image, b) vertical flipping, c) random rotation.

**Supplementary Table 1.** Adjusted (Bonferroni correction) p-values for the pairwise t-tests between average spectra for each disease, or pair of diseases, and uninoculated leaves, for each of the 19 wavelengths. p-values corresponding to the four largest tests statistics for each disease class are highlighted in bold.

| Wavelength | Mildew | Yellow rust | Septoria | YR + septoria | YR + mildew |
| --- | --- | --- | --- | --- | --- |
| **375** | $3.14 \times 10^{-4}$ | $2.14 \times 10^{-30}$ | $6.74 \times 10^{-9}$ | $3.05 \times 10^{-34}$ | $7.70 \times 10^{-23}$ |
| **405** | $4.45 \times 10^{-4}$ | $1.78 \times 10^{-104}$ | $2.96 \times 10^{-19}$ | $2.95 \times 10^{-11}$ | $2.11 \times 10^{-20}$ |
| **435** | $1.22 \times 10^{-6}$ | $7.63 \times 10^{-109}$ | $2.60 \times 10^{-9}$ | $4.11 \times 10^{-53}$ | $1.20 \times 10^{-10}$ |
| **450** | $8.88 \times 10^{-6}$ | $2.05 \times 10^{-123}$ | $9.58 \times 10^{-130}$ | $1.46 \times 10^{-151}$ | $7.87 \times 10^{-23}$ |
| **470** | $6.81 \times 10^{-2}$ | $1.26 \times 10^{-148}$ | $7.32 \times 10^{-180}$ | $4.80 \times 10^{-197}$ | $1.25 \times 10^{-38}$ |
| **505** | 1 | $1.47 \times 10^{-191}$ | $2.38 \times 10^{-204}$ | $7.61 \times 10^{-223}$ | $2.79 \times 10^{-62}$ |
| **525** | $4.73 \times 10^{-5}$ | $4.11 \times 10^{-160}$ | $3.90 \times 10^{-52}$ | $1.41 \times 10^{-62}$ | $2.26 \times 10^{-8}$ |
| **570** | $9.94 \times 10^{-4}$ | $2.43 \times 10^{-177}$ | $1.26 \times 10^{-67}$ | $2.36 \times 10^{-91}$ | $1.34 \times 10^{-24}$ |
| **590** | $2.83 \times 10^{-6}$ | $3.87 \times 10^{-279}$ | $4.19 \times 10^{-113}$ | $1.67 \times 10^{-143}$ | $6.21 \times 10^{-30}$ |
| **630** | $5.84 \times 10^{-5}$ | 0.00 | $2.52 \times 10^{-244}$ | $8.58 \times 10^{-260}$ | $5.92 \times 10^{-48}$ |
| **645** | $1.69 \times 10^{-4}$ | **0.00** | **0.00** | **0.00** | $2.04 \times 10^{-66}$ |
| **660** | $6.31 \times 10^{-3}$ | **0.00** | **0.00** | **0.00** | **$4.67 \times 10^{-101}$** |
| **700** | $5.33 \times 10^{-4}$ | **0.00** | **0.00** | **0.00** | **$2.85 \times 10^{-97}$** |
| **780** | $1.38 \times 10^{-3}$ | **0.00** | **0.00** | **0.00** | $1.44 \times 10^{-85}$ |
| **850** | $2.21 \times 10^{-12}$ | $3.75 \times 10^{-39}$ | $2.26 \times 10^{-87}$ | $9.42 \times 10^{-75}$ | $1.27 \times 10^{-39}$ |
| **870** | **$7.56 \times 10^{-14}$** | $4.67 \times 10^{-53}$ | $8.46 \times 10^{-34}$ | $5.24 \times 10^{-88}$ | $1.37 \times 10^{-69}$ |
| **890** | **$2.54 \times 10^{-15}$** | $6.30 \times 10^{-59}$ | $9.02 \times 10^{-102}$ | $2.67 \times 10^{-115}$ | $1.69 \times 10^{-96}$ |
| **940** | **$1.31 \times 10^{-16}$** | $1.39 \times 10^{-91}$ | $1.43 \times 10^{-39}$ | $2.21 \times 10^{-198}$ | **$1.11 \times 10^{-155}$** |
| **970** | **$3.87 \times 10^{-17}$** | $4.61 \times 10^{-143}$ | $8.44 \times 10^{-42}$ | $4.95 \times 10^{-245}$ | **$6.24 \times 10^{-185}$** |

**Supplementary Table 2.** Adjusted (Bonferroni correction) p-values for the pairwise t-tests between average spectra for the individual diseases and combinations of diseases containing them, for each of the 19 wavelengths. p-values significant at 5% are highlighted in bold.

| Wavelength | Mildew vs<br>YR + mildew | Yellow rust vs<br>YR + mildew | Septoria vs<br>YR + septoria | Yellow rust vs<br>YR + septoria |
| --- | --- | --- | --- | --- |
| 375 | **6.20 x 10<sup>-4</sup>** | 1 | **1.70 x 10<sup>-11</sup>** | 1 |
| 405 | 2.54 x 10<sup>-1</sup> | **106 x 10<sup>-6</sup>** | **5.46 x 10<sup>-4</sup>** | **6.58 x 10<sup>-5</sup>** |
| 435 | 1.09 x 10<sup>-1</sup> | **4.19 x 10<sup>-7</sup>** | **8.39 x 10<sup>-2</sup>** | 6.55 x 10<sup>-1</sup> |
| 450 | 5.46 x 10<sup>-1</sup> | **2.96 x 10<sup>-5</sup>** | 1 | 1 |
| 470 | 1 | **1.66 x 10<sup>-4</sup>** | 1 | 3.64 x 10<sup>-1</sup> |
| 505 | 1 | **6.95 x 10<sup>-5</sup>** | 1 | 9.83 x 10<sup>-1</sup> |
| 525 | 1 | **2.71 x 10<sup>-14</sup>** | 1 | **4.38 x 10<sup>-11</sup>** |
| 570 | 1 | **4.70 x 10<sup>-13</sup>** | 1 | **8.18 x 10<sup>-10</sup>** |
| 590 | 1.19 x 10<sup>-1</sup> | **5.49 x 10<sup>-19</sup>** | 1 | **5.68 x 10<sup>-22</sup>** |
| 630 | 1.02 x 10<sup>-1</sup> | **8.21 x 10<sup>-18</sup>** | 1 | **9.22 x 10<sup>-15</sup>** |
| 645 | **4.79 x 10<sup>-2</sup>** | **3.21 x 10<sup>-16</sup>** | **1.90 x 10<sup>-2</sup>** | **1.48 x 10<sup>-5</sup>** |
| 660 | **3.64 x 10<sup>-2</sup>** | **5.33 x 10<sup>-15</sup>** | **6.80 x 10<sup>-4</sup>** | 2.52 x 10<sup>-1</sup> |
| 700 | **1.05 x 10<sup>-2</sup>** | **1.77 x 10<sup>-14</sup>** | **9.92 x 10<sup>-6</sup>** | 1 |
| 780 | **7.58 x 10<sup>-2</sup>** | **5.04 x 10<sup>-15</sup>** | **1.27 x 10<sup>-3</sup>** | 1 |
| 850 | **2.16 x 10<sup>-2</sup>** | 1 | 1 | 1 |
| 870 | **1.72 x 10<sup>-2</sup>** | 3.38 x 10<sup>-1</sup> | 1.49 x 10<sup>-1</sup> | 1 |
| 890 | **1.24 x 10<sup>-2</sup>** | **2.35 x 10<sup>-4</sup>** | 1 | 1 |
| 940 | **9.30 x 10<sup>-2</sup>** | **1.37 x 10<sup>-6</sup>** | **1.40 x 10<sup>-11</sup>** | **4.05 x 10<sup>-3</sup>** |
| 970 | 2.93 x 10<sup>-1</sup> | **2.36 x 10<sup>-6</sup>** | **2.34 x 10<sup>-17</sup>** | **1.47 x 10<sup>-2</sup>** |

## References

1. Food and Agriculture Organization of the United Nations [Internet]. [cited 2025 Aug 21]. Available from: https://www.fao.org/faostat/en/#data/QCL

2. Figueroa M, Hammond-Kosack KE, Solomon PS. A review of wheat diseases-a field perspective: A review of wheat diseases. Mol Plant Pathol. 2018;19(6):1523–36.

3. Genaev MA, Skolotneva ES, Gultyaeva EI, Orlova EA, Bechtold NP, Afonnikov DA. Image-Based Wheat Fungi Diseases Identification by Deep Learning. Plants. 2021;10(8)1500.

4. Chen W, Wellings C, Chen X, Kang Z, Liu T. Wheat stripe (yellow) rust caused by Puccinia striiformis f. sp. tritici. Mol Plant Pathol. 2014;15(5):433–46.

5. Smith HC, Blair ID. Wheat powdery mildew investigations. Ann Appl Biol. 1950;37(4):570–83.

6. Orton ES, Brown JK. Reduction of growth and reproduction of the biotrophic fungus Blumeria graminis in the presence of a necrotrophic pathogen. Front Plant Sci. 2016;7:742.

7. Barbedo JGA. A review on the main challenges in automatic plant disease identification based on visible range images. Biosyst Eng. 2016 Apr 1;144:52–60.

8. Dutt A, Andrivon D, Le May C. Multi-infections, competitive interactions, and pathogen coexistence. Plant Pathol. 2022;71(1):5–22.

9. Bock CH, Poole GH, Parker PE, Gottwald TR. Plant Disease Severity Estimated Visually, by Digital Photography and Image Analysis, and by Hyperspectral Imaging. Crit Rev Plant Sci. 2010;29(2):59–107.

10. Farber C, Mahnke M, Sanchez L, Kurouski D. Advanced spectroscopic techniques for plant disease diagnostics. A review. TrAC Trends Anal Chem. 2019;118:43–9.

11. Golhani K, Balasundram SK, Vadamalai G, Pradhan B. A review of neural networks in plant disease detection using hyperspectral data. Inf Process Agric. 2018;5(3):354–71.

12. Lowe A, Harrison N, French AP. Hyperspectral image analysis techniques for the detection and classification of the early onset of plant disease and stress. Plant Methods. 2017;13:80.

13. Mishra P, Asaari MSM, Herrero-Langreo A, Lohumi S, Diezma B, Scheunders P. Close range hyperspectral imaging of plants: A review. Biosyst Eng. 2017;164:49–67.

14. Gomes C, Costa R, Almeida A, Coutinho J, Pinheiro N, Coco J, et al. Septoria leaf blotch and yellow rust control by: fungicide application opportunity and genetic response of bread wheat varieties. Emir J Food Agric. 2016;28(7): 493–500.

15. Spadafora V, Cole Jr H. Interactions between Septoria nodorum leaf blotch and leaf rust on soft red winter wheat. Phytopathology.1987;77(9):1308–10.

16. Van der Wal A, Shearer B, Zadoks J. Interaction between Puccinia recondita f. sp. triticina and Septoria nodorum on wheat, and its effects on yield. Neth J Plant Pathol. 1970;76(4):261–3.

17. Madariaga R, Scharen, A. L. Interactions of Puccinia striiformis and Mycosphaerella graminicola on Wheat. Plant Dis. 1986;70(7):651.

18. Kamilaris A, Prenafeta-Boldú FX. Deep learning in agriculture: A survey. Comput Electron Agric. 2018;147:70–90.

19. Nagaraju M, Chawla P. Systematic review of deep learning techniques in plant disease detection. Int J Syst Assur Eng Manag. 2020;11(3):547–60.

20. Saleem MH, Potgieter J, Arif KM. Plant Disease Detection and Classification by Deep Learning. Plants. 2019;8(11):468.

21. Boulent J, Foucher S, Théau J, St-Charles PL. Convolutional Neural Networks for the Automatic Identification of Plant Diseases. Front Plant Sci. 2019;10:941.

22. Guo A, Huang W, Ye H, Dong Y, Ma H, Ren Y, et al. Identification of Wheat Yellow Rust Using Spectral and Texture Features of Hyperspectral Images. Remote Sens. 2020;12(9):1419.

23. Cheshkova AF. A review of hyperspectral image analysis techniques for plant disease detection and identif ication. Vavilov J Genet Breed. 2022;26(2):202–13.

24. Oppelt N, Mauser W. Hyperspectral monitoring of physiological parameters of wheat during a vegetation period using AVIS data. Int J Remote Sens. 2004;25(1):145–59.

25. De Bei R, Cozzolino D, Sullivan W, Cynkar W, Fuentes S, Dambergs R, et al. Non-destructive measurement of grapevine water potential using near infrared spectroscopy. Aust J Grape Wine Res. 2011;17(1):62–71.

26. Szegedy C, Liu W, Jia Y, Sermanet P, Reed S, Anguelov D, et al. Going deeper with convolutions. In: Proceedings of the IEEE conference on computer vision and pattern recognition. 2015:1–9.

27. Tan M, Le QV. EfficientNet: Rethinking Model Scaling for Convolutional Neural Networks. In: International conference on machine learning. 2019:6105–14.

28. Nass HG, Pridham ED, Mellish D, Langille JE, Walker DW. Vuka winter wheat. Can J Plant Sci. 1985;65(4):1083–4.

29. VideometerLab 4 Technical Specifications [Internet]. [cited 2025 Aug 21]. Available from: https://videometer.com/wp-content/uploads/2021/12/VideometerLab_2021_WithoutCrop.pdf

30. Qi C, Sandroni M, Cairo Westergaard J, Høegh Riis Sundmark E, Bagge M, Alexandersson E, et al. In-field classification of the asymptomatic biotrophic phase of potato late blight based on deep learning and proximal hyperspectral imaging. Comput Electron Agric. 2023;205:107585.

31. Yu S, Jia S, Xu C. Convolutional neural networks for hyperspectral image classification. Neurocomputing. 2017;219:88–98.

32. Nguyen C, Sagan V, Maimaitiyiming M, Maimaitijiang M, Bhadra S, Kwasniewski MT. Early Detection of Plant Viral Disease Using Hyperspectral Imaging and Deep Learning. Sensors. 2021;21(3):742.

33. Lu Y, Chen D, Olaniyi E, Huang Y. Generative adversarial networks (GANs) for image augmentation in agriculture: A systematic review. Comput Electron Agric. 2022;200:107208.

34. Egusquiza I, Benito-Del-Valle L, Picón A, Bereciartua-Pérez A, Gómez-Zamanillo L, Elola A, et al. When synthetic plants get sick: Disease graded image datasets by novel regression-conditional diffusion models. Comput Electron Agric. 2025;229:109690.

35. Xie C, Shao Y, Li X, He Y. Detection of early blight and late blight diseases on tomato leaves using hyperspectral imaging. Sci Rep. 2015;5(1):16564.

36. Hsiao CF, Feyrer G, Stein A. Classifying early-stage soybean fungal diseases on hyperspectral images using convolutional neural networks. Smart Agric Technol. 2025;11:101023.

37. Dutta A, Tyagi R, Chattopadhyay A, Chatterjee D, Sarkar A, Lall B, et al. Early detection of wilt in Cajanus cajan using satellite hyperspectral images: Development and validation of disease-specific spectral index with integrated methodology. Comput Electron Agric. 2024;219:108784.

38. Matese A, Czarnecki JMP, Samiappan S, Moorhead R. Are unmanned aerial vehicle-based hyperspectral imaging and machine learning advancing crop science? Trends Plant Sci. 2024;29(2):196–209.

39. Abdulridha J, Min A, Rouse MN, Kianian S, Isler V, Yang C. Evaluation of Stem Rust Disease in Wheat Fields by Drone Hyperspectral Imaging. Sensors. 2023;23(8):4154.

40. Madariaga RB. Interactions of Puccinia striiformis and Mycosphaerella graminicola on wheat [PhD Thesis]. Montana State University-Bozeman, College of Agriculture; 1984.

41. Zhou C, Zhao Z, Chen W, Feng Y, Song J, Xiang W. Residual attention based multi-label learning for apple leaf disease identification.J Agric Eng. 2024;55(4).

42. Li H, Huang L, Ruan C, Huang W, Wang C, Zhao J. A dual-branch neural network for crop disease recognition by integrating frequency domain and spatial domain information. Comput Electron Agric. 2024;219:108843.

43. Deng J, Dong W, Socher R, Li LJ, Li K, Fei-Fei L. ImageNet: A large-scale hierarchical image database. In: 2009 IEEE Conference on Computer Vision and Pattern Recognition. 2009:248–55.

44. Xie C, He Y. Spectrum and Image Texture Features Analysis for Early Blight Disease Detection on Eggplant Leaves. Sensors. 2016;16(5):676.

45. Selvaraju RR, Cogswell M, Das A, Vedantam R, Parikh D, Batra D. Grad-CAM: Visual Explanations from Deep Networks via Gradient-Based Localization. In: 2017 IEEE International Conference on Computer Vision (ICCV). 2017:618–26.

46. Chicco D. Siamese Neural Networks: An Overview. Methods Mol Biol Clifton NJ. 2021;2190:73–94.

47. Sung F, Yang Y, Zhang L, Xiang T, Torr PHS, Hospedales TM. Learning to Compare: Relation Network for Few-Shot Learning. In: Proceedings of the IEEE conference on computer vision and pattern recognition. 2018:1199–208.

48. Chollet F. Keras [Internet]. 2015. Available from: https://keras.io

